# Assessing the acoustic behaviour of *Anopheles gambiae* s.l. *dsxF* mutants: Implications for Vector Control

**DOI:** 10.1101/2020.09.06.284679

**Authors:** Matthew P Su, Marcos Georgiades, Judit Bagi, Kyros Kyrou, Andrea Crisanti, Joerg T Albert

**Author notes:** These authors contributed equally (Matthew P Su, Marcos Georgiades). E-mails: MS MG JB KK AC JTA.

## Abstract

**Background:** The release of genetically modified mosquitoes which use gene-drive mechanisms to suppress reproduction in natural populations of *Anopheles* mosquitoes is one of the scientifically most promising methods for malaria transmission control. However, many scientific, regulatory and ethical questions remain before transgenic mosquitoes can be utilised in the field. Mutations which reduce an individual’s reproductive success are likely to create strong selective pressures to evolve resistance. It is thus crucial that the targeted population collapses as rapidly and as completely as possible to reduce the available time for the emergence of drive-resistant mutations. At a behavioural level, this means that the gene-drive carrying mutants should be at least as (and ideally more) sexually attractive than the wildtype population they compete against. A key element in the copulatory negotiations of *Anopheles* mosquitoes is their acoustic courtship. We therefore analysed sound emissions and acoustic preference in a *doublesex* mutant previously used to successfully collapse caged colonies of *Anopheles gambiae s*.*l*..

**Methods:** The flight tones produced by the beating of their wings form the signals for acoustic mating communication in *Anopheles* species. We assessed the acoustic impact of the disruption of a female-specific isoform of the *doublesex* gene (*dsxF*) on the wing beat frequency (WBF; measured as *flight tone*) of both males (XY) and females (XX) in homozygous *dsxF*^*-*^ mutants (*dsxF*^*-/-*^), heterozygous *dsxF*^*-*^ carriers (*dsxF*^*+/-*^) and G3 ‘wildtype’ *dsxF*^*+*^ controls (*dsxF*^*+/+*^). To exclude non-genetic influences, we controlled for temperature and measured wing lengths for all experimental animals. We used a phonotaxis assay to test the acoustic preferences of mutant and control mosquitoes.

**Results:** A previous study demonstrated an altered phenotype only for females homozygous for the disrupted *dsx* allele (*dsxF*^*-/-*^), who appear intersex. No phenotypic changes were observed for heterozygous carriers or males, suggesting that the female-specific *dsxF* allele is haplosufficient. We here identify significant, dose-dependent increases in the flight tones of both *dsxF*^*-/-*^ and *dsxF*^*+/-*^ females when compared to *dsxF*^*+/+*^ control females. Flight tone frequencies in all three female genotypes remained significantly lower than in males, however. When tested experimentally, males showed stronger phonotactic responses to the flight tones of control *dsxF*^*+/+*^ females. While flight tones from *dsxF*^*+/-*^ and *dsxF*^*-/-*^ females also elicited positive phonotactic behaviour in males, this was significantly reduced compared to responses to control tones. We found no evidence of phonotactic behaviour in any female genotype tested. None of the male genotypes displayed any deviations from the control condition.

**Conclusions:** A key prerequisite for copulation in anopheline mosquitoes is the phonotactic attraction of males towards female flight tones within large - spatially and acoustically crowded - mating swarms. Reductions in acoustic attractiveness of released mutant lines, as reported here for heterozygous *dsxF*^*+/-*^ females, reduce the line’s mating efficiency, and could consequently reduce the efficacy of the associated population control effort. Assessments of caged populations may not successfully reproduce the challenges posed by natural mating scenarios. We propose to amend existing testing protocols in order to more faithfully reflect the competitive conditions between a mutant line and the wildtype population it is meant to interact with. This should also include novel tests of ‘acoustic fitness’. In line with previous studies, our findings confirm that disruption of the female-specific isoform *dsxF* has no effect on males; for some phenotypic traits, such as female flight tones, however, the effects of *dsxF* appear to be dose-dependent rather than haplosufficient.

## Background

Mosquitoes represent a major global health problem, with *Aedes, Anopheles* and *Culex* species acting as vectors of diseases that infect millions of people each year [1]. Malaria remains a major cause of mortality and morbidity worldwide in spite of significant advances made in disease control since the turn of the century [2, 3]. This is in part due to the reduced efficacy of current control tools such as insecticidal nets and indoor residual spraying, as well as the emergence of secondary disease vectors [4, 5, 6]. Novel control techniques are therefore necessary to continue the push towards disease elimination [7].

One potential option is the utilisation of gene drive systems, which target haplosufficient female fertility genes, leading to a reduction in female fertility and, eventually, population collapse [8, 9]. The recent generation of *Anopheles gambiae* CRISPR/Cas9 mutants in which a female specific exon of the *doublesex* (*dsxF*) gene was disrupted is here of interest. Lab cage trials have demonstrated that the introduction of *dsxF* mutants into cages of wildtype mosquitoes was sufficient to lead to eventual population collapse [10].

However, there are many scientific, ethical and regulatory hurdles to overcome before such transgenic mosquitoes can be released in even semi-field trials [11]. It is vital that any transgenic mosquitoes are subjected to rigorous testing prior to use in the field; gene transfer into natural populations following release of transgenic *Aedes aegypti* has highlighted the potential risks of release of transgenic insects [12]. On a scientific level, one important task will be to maintain the gene drive’s effectiveness outside of the laboratory and under more ‘real world’ scenarios.

A major element of this testing is the investigation of interactions with natural, non-mutant populations, particularly with regards to courtship behaviour. If mutant mosquitoes are unable, or only less likely, to copulate with native populations then they become the less attractive option, which will slow down or outright frustrate the population control effort [13]. In addition to potential direct and indirect fitness costs associated with mutations, laboratory habituation and mass rearing can also affect mating performance [14, 15]. In this context it is noteworthy that the *dsxF* mutants we tested were also generated from a lab-established strain (G3) rather than any wildtype population [10]. Extensive testing of mutant mating fitness prior to translation from laboratory mating assays is thus a key requirement for assessing a specific line’s suitability for use as part of a release program.

The sense of hearing is a vital component of mosquito reproduction, with males identifying females within swarms via phonotactic responses to female flight tones and acoustic communication is also thought to play a role in female mate selection [16, 17, 18, 19]. The phonotactic response is highly specific, however, with males responding only to a narrow range of frequencies [20]. Both male and female mosquitoes have extraordinarily sensitive and complex ears, but there are also significant sexual dimorphisms in auditory function and hearing-related behaviours [21, 22, 23].

Chromosomally female (XX) *dsxF*^*-/-*^ mutants display an intersex phenotype, which also includes an intersex morphology of their flagellar sound receivers [10]; If, and if so to what extent, other parts of the auditory or acoustic system are affected by the allelic disruption is unclear. Physiological changes that could impact the mutants’ ability to interbreed with existing mosquitoes, are e.g. changes in male or female flight tones – or their corresponding acoustic preferences. It is currently unclear if any of the *dsxF* mutant genotypes affects these parameters. If so, this could have substantial effects on the ability of mutants to interbreed with existing mosquitoes.

In order to address this topic, we tested the flight tones and phonotactic responses of *dsxF* XX and XY mutants and controls. We found that whilst male (XY) mutant (*dsxF*^*-/-*^, *dsxF*^*+/-*^) flight tones were not significantly different to male controls (*dsxF*^*+/+*^), female (XX) mutant (*dsxF*^*-/-*^, *dsxF*^*+/-*^) flight tones had significantly higher frequencies than those of their respective controls (*dsxF*^*+/+*^), with both showing an increase towards the male flight tone in a seemingly dose-response fashion.

No female showed evidence of phonotaxis to any of the acoustic stimuli we provided, whilst all males showed a strong phonotactic response to tones of 400Hz (but much reduced or absent responses to tones of 100Hz or 700Hz). However, a more focused phonotaxis assay using the median flight tones obtained from each of the three female genotypes (*dsxF*^*+/+*^, *dsxF*^*+/-*^, *dsxF*^*-/-*^) found that control males responded far more strongly to the flight tones of control females than to either of the mutant flight tones. Preliminary tests of *dsxF*^*-/-*^ males showed a similar preference for control flight tones (Supplemental Figure 2). As such, it seems likely that male mosquitoes of any genotype will demonstrate a strong preference for wildtype females, with mutant females potentially reduced to a lesser attractive role.

**Figure 1:**
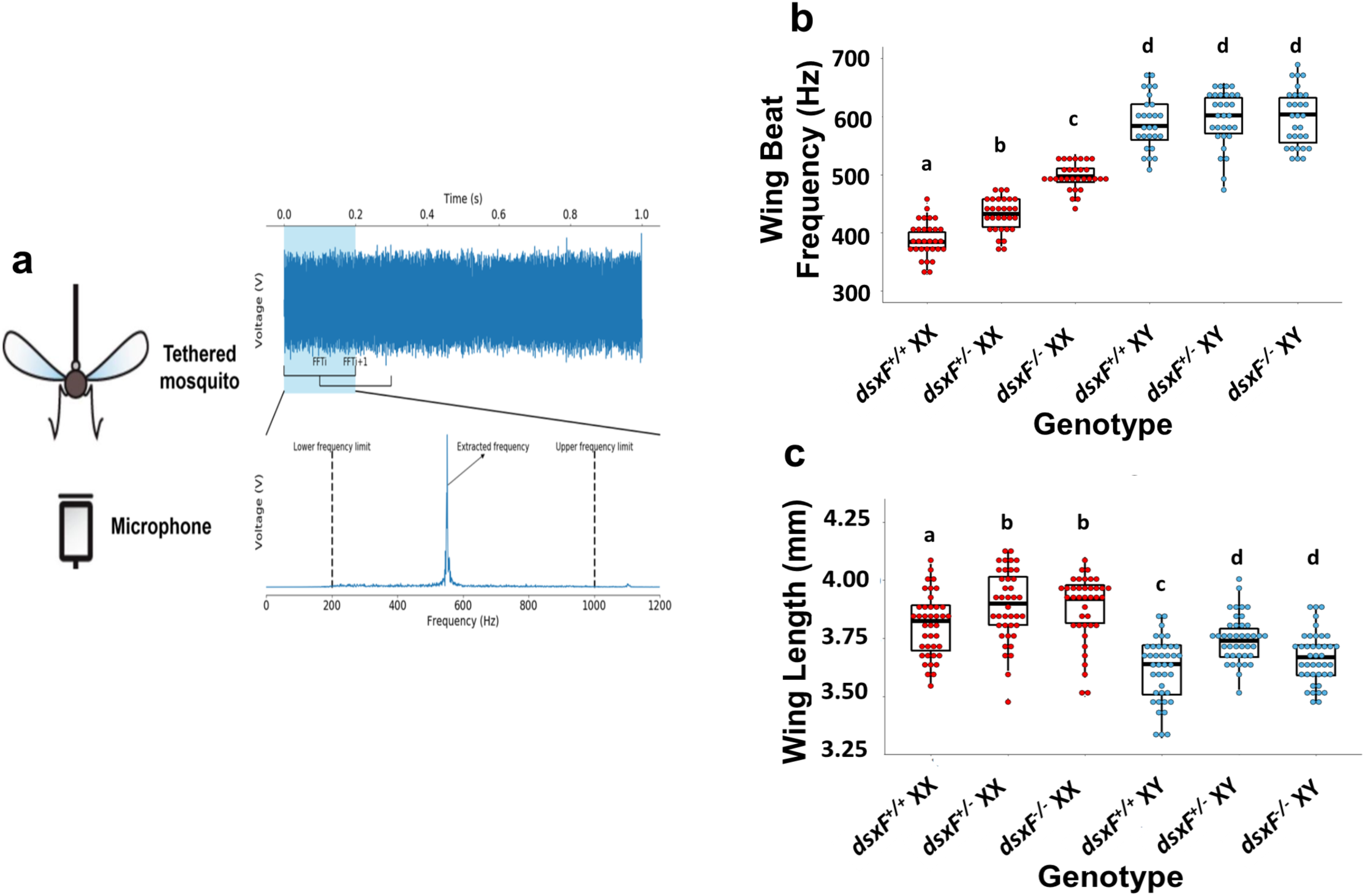
dsxF^+/-^and dsxF^-/-^ XX mutants have different wing beat frequencies (=flight tones) to all other groups. a) Sketch of flight tone recording set-up: mosquitoes were tethered then placed at a constant distance from a microphone. Temperature and humidity conditions were controlled (21-22°C; 50% RH) and recordings always took place within the same two-hour window. b) Calculated wing beat frequencies for each genotype – significant differences (Two-way *ANOVA; * p < 0*.*05*) between groups are indicated by letter. Centre line mean; box limits, lower and upper quartiles; whiskers, 5th and 95th percentiles (identical B-C). Sample sizes: *dsxF*^*+/+*^ XX = 30; *dsxF*^*+/-*^ XX = 30; *dsxF*^*-/-*^ XX = 30; *dsxF*^*+/+*^ XY = 27; *dsxF*^*+/-*^ XY = 30; *dsxF*^*-/-*^ XY = 30. c) Wing length measurements for each genotype - significant differences (Two-way *ANOVA; * p < 0*.*05*) between groups are indicated by letter. Sample sizes: *dsxF*^*+/+*^ XX = 40; *dsxF*^*+/-*^ XX = 40; *dsxF*^*-/-*^ XX = 40; *dsxF*^*+/+*^ XY = 41; *dsxF*^*+/-*^ XY = 40; *dsxF*^*-/-*^ XY = 41.

**Figure 2:**
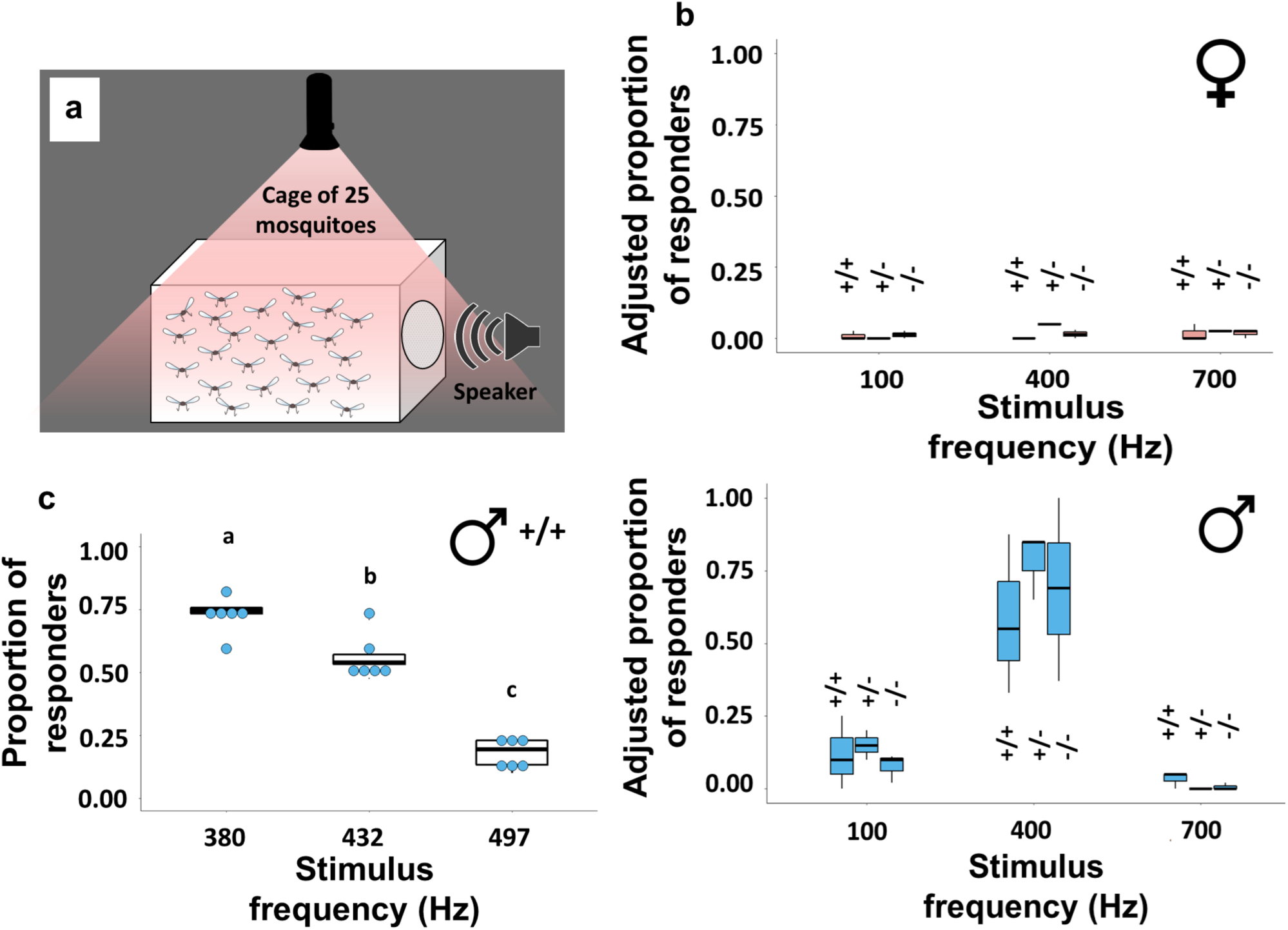
Males show a strong preference for acoustic stimuli of similar frequency to wildtype female flight tones; this phonotactic response is reduced as the tone becomes increasingly different. a) Diagram of phonotaxis experimental set-up: Single-sex virgin cages were provided with one-minute periods of stimulation in the form of three pure tones (100, 400 and 700Hz) or a one-minute period of silence. The number of mosquitoes attracted to the sound source for each type of stimulus was calculated. b) Adjusted proportion of mosquitoes responding to each stimulus type (no stimulus, 100Hz, 400Hz and 700Hz respectively) for XX and XY mosquitoes from each genotype. Centre circle, median; error bars represent ± SEM. c) Adjusted proportion of control mosquitoes responding to each stimulus type (380Hz, 432Hz and 497Hz respectively) for *dsxF*^*+/+*^ XY mosquitoes. Centre line, median; error bars represent ± SEM.

## Methods

### Mosquito rearing

*An. gambiae* G3 strain (*dsxF*^+/+^), as well as *dsxF*^+/-^ and *dsxF*^-/-^ mutant pupae, were reared and provided by the Crisanti lab at Imperial College London. Larval density was kept constant throughout the rearing process.

*dsxF*^+/+^ and *dsxF*^+/-^ pupae were sex separated and kept in single sex cages in incubators maintained at 28°C and 80% relative humidity. Light/ dark conditions included a one-hour ramping period of constantly increasing white light; ramp for lights-ON from zeitgeber time (ZT) ZT0 to ZT1; then 11 hours (ZT1-ZT12) of white light at constant intensity, followed by a one-hour (ZT12-13) ramping period of constantly decreasing white light, and then 11 hours (ZT13-ZT24) of constant darkness. All light ramps transitioned linearly between a Photosynthetic Photon Flux Density (PPFD) of 80 µmol/m^2^/s (or ∼ 5929 lux) and complete darkness (0 µmol/m^2^/s or 0 lux, respectively).

*dsxF*^*-/-*^ pupae were not sex separated but were otherwise reared in identical conditions. Mosquitoes were supplied with a constant source of 10% glucose solution.

All mosquitoes used for experiments were virgin and aged 3 – 7 days old. Mosquitoes were housed within temperature, humidity and light-controlled incubators for three days prior to all experiments, which were conducted during a time corresponding to sunset (which represents a time period of peak activity and swarming under natural conditions).

### Wing length measurements

The right wings of adult mosquitoes from each genotype were removed using a pair of forceps whilst the mosquitoes were CO2 sedated. The wings and flagellae were then transferred to separate microscope slides in groups of five. Each individual sample was immediately imaged using a Zeiss Axioplan 2 microscope and Axiovision 4.3 software. Wing lengths were determined using the Axiovision 4.3 software length measurement function, calibrated to the nearest 0.1 mm. Three biological repeats were conducted over separate generations.

Total sample sizes for each group: *dsxF*^*+/+*^ XX = 40; *dsxF*^*+/-*^ XX = 40; *dsxF*^*-/-*^ XX = 40; *dsxF*^*+/+*^ XY = 41; *dsxF*^*+/-*^ XY = 40; *dsxF*^*-/-*^ XY = 41.

### Wing beat frequency measurements

A resin casing was printed using an Ultimaker 2+ 3D printer and used to house a particle velocity microphone (Knowles NR-3158). The whole apparatus was held in a micromanipulator placed on a vibration isolation table.

Adult mosquitoes from each genotype were cold-sedated using ice before blue-light cured glue was used to fix the tip of a tungsten wire to their thoraces, taking care not to restrict or damage the wings in doing so. The tethered mosquito was mounted into the microphone case and oriented such that its posterior was facing the particle velocity microphone. All measurements were conducted in the same isolated room at a temperature between 21 – 22°C.

Mosquito flight was initiated via a tarsal reflex response [24]. A small cotton ball was placed underneath each tethered mosquito; once the mosquito had clasped the ball, it was swiftly removed, with this removal stimulating flight initiation. Minimum flight length used was 10 seconds. The voltage timeseries waveform measured for each flying mosquito by the particle velocity microphone was recorded using the Spike2 software (Cambridge Electronic Design Ltd., UK).

Sample sizes for each group were: *dsxF*^*+/+*^ XX = 30; *dsxF*^*+/-*^ XX = 30; *dsxF*^*-/-*^ XX = 30; *dsxF*^*+/+*^ XY = 27; *dsxF*^*+/-*^ XY = 30; *dsxF*^*-/-*^ XY = 30.

### Flight tone analysis algorithm

Raw data from the Spike2 recordings were exported to Python for analysis via a custom script. The first and last two seconds of each flight were discarded prior to analysis. Subsequently the timeseries was divided into 5-second subsegments, discarding the final shorter subsegment (if flight length modulo 5 ≠ 0). A Fast Fourier transform (FFT) with a 200ms window was then applied throughout each of the subsegments. This window was shifted in 100ms increments

(i.e. 50% overlap between successive FFTs) and applied repeatedly until the end of the flight segment was reached (Figure 1a).

Limits were applied to the frequency domain of each FFT such that only frequencies between 200 – 1,000Hz would be extracted for analysis. For each FFT, the peak frequency was identified and assigned as the flight tone for the time segment over which the FFT was calculated. A list of peak frequencies was compiled for each of the aforementioned subsegments. These lists were added together and averaged, resulting in a 5-second long final list of average frequencies. The mean was employed in the averaging step as these values were normally distributed. As the list of means, in turn, tended to be non-normally distributed, the median was taken and assigned as the flight tone of that individual animal. The segmentation of the original waveform and summarization into a single 5-second long list of values served to moderate for potential effects of flight duration on the animals’ flight tone.

### Spectrally broad phonotaxis assay

Female (XX) and male (XY) mosquitoes from all three genotypes were aspirated into small, single-sex cages in groups of 25 and kept for at least two hours in the same room used for flight tone experiments. All experiments were conducted at a temperature between 20 – 23°C and at ∼ ZT13 (i.e. swarming time, around the time of complete cessation of light). Throughout the experiment, mosquitoes were kept in constant darkness.

A free app (TMsoft tone generator) was used to provide acoustic stimulation to caged mosquitoes; this stimulation consisted of three pure tones with frequencies of 100, 400 and 700 Hz. These frequencies were chosen based on the prior recordings which found no female flight tones as high as 700Hz, and no male or female frequency as low as 100Hz.

The sound source was placed next to the cage with its speaker touching the cage mesh prior to stimulus initiation. Each tone was played for 1 minute and was succeeded in turn by a 1-minute long silence before the next tone was played. The tones were played first from low to high frequencies and subsequently from high to low, allowing mosquitoes to rest for 5 minutes between forward and backward playbacks. To ensure that mosquitoes were being attracted to the sound emitted by the sound source rather than the sound source itself, at the start of each experiment the sound source was placed next to the cage with its speaker touching the cage’s mesh with no stimulus playing. Mosquitoes that approached the sound source during either control or acoustic playback were counted manually using a red-light flashlight. Three biological repeats were conducted for each group.

### Spectrally focused phonotaxis assay

*dsxF*^*+/+*^ XY mosquitoes were tested in groups of 25 as above for the broad-range assay, this time, however using three pure tones with frequencies equal to the recorded median flight tone frequencies of each of the female genotypes; 380Hz (*dsxF*^*+/+*^), 432Hz (*dsxF*^*+/-*^) and 497Hz (*dsxF*^*-/-*^). [Please note that the played *dsxF*^*+/+*^ control tone of 380Hz is marginally different from the median reported in Table 1; this is because the flight tone choice for the playback experiments was based on an earlier data cohort].

**Table 1:**
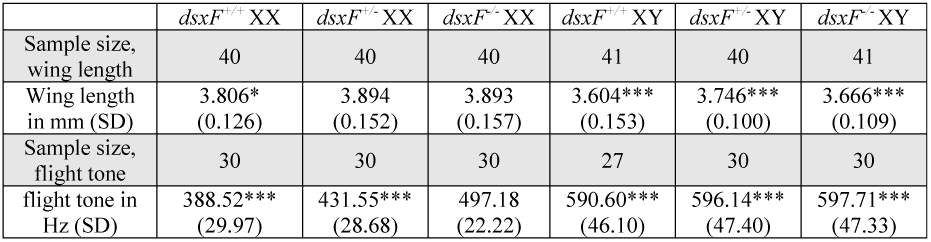
Quantification of changes to dsxF^+/-^ XX flight tones. Mean values of wing lengths and flight tones for dsxF^+/+^, dsxF^+/-^ and dsxF^+/+^ XX and XY mosquitoes, with standard deviation (SD) values provided in brackets. Significant differences found between dsxF^-/-^ XX mosquitoes and any other mosquito group are starred (ANOVA on ranks; *p < 0.05; ***p < 0.001).

## Statistical analysis

Flight tone analyses were conducted in Python. Remaining analyses were completed in Matlab and R. Throughout the analyses, all statistical tests used a significance level of p <0.05.

Sample sizes for all experiments were determined via reference to published investigations. Within-group variation estimates were calculated when appropriate as part of standard statistical testing.

Statistical tests for normality (Shapiro–Wilk Normality tests with a significance level of p < 0.05) were first applied to each dataset.

Wing length measurements and flight tones were found to be normally distributed; two-way ANOVA tests were thus used for comparisons across the genotypes and sexes.

For the spectrally broad phonotaxis assay, the proportion of responders to the control stimulus (silence) was subtracted from the proportion of responders to the stimulus tones. That is, to calculate the adjusted proportion of responders we calculated:

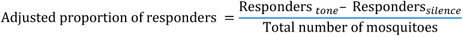

Where Responders_tone_ and Responders_silence_ refers to the number of responders to the individual tones or silence respectively. One-way ANOVAs were then used to test for differences in responses between the stimulus tone frequencies. For the focused phonotaxis assay, no adjusted proportion was calculated and one-way ANOVAs were applied directly to the Proportion of responders.

## Results

### *dsxF+/-* and *dsxF-/-* XX mutants have different flight tones to all other XX and XY mosquitoes

By recording the flight tones of tethered female and male mosquitoes we were able to calculate the median flight tones for each group (Figure 1b). All male flight tones were found to be greater than all female flight tones, but we found no differences between males (Two-way ANOVA; p<0.001; p>0.05 respectively). The flight tones of *dsxF*^*-/-*^ females were significantly different from all other groups; they were significantly higher than the other female genotypes (497 ± 22.2 Hz compared to 432 ± 28.7 Hz and 380 ± 30.0 Hz for *dsxF*^*+/-*^ and *dsxF*^*+/+*^, respectively), and significantly lower than all male genotypes (Two-way ANOVA; p<0.001; Table 1; Figure 1b). We also found a significant difference between *dsxF*^*+/-*^ XX mutants and the other two female genotypes in an apparent dose response fashion (Two-way ANOVA; p=0.002; Figure 1b).

Mosquito flight tone frequencies have been reported to show correlations with temperature (see e.g. [25] for *Aedes*), but the relation, especially for anopheline mosquitoes, has remained unclear. Here, temperature was tightly controlled, with all recordings being made between 21 and 22^°^C. The relationship between wing beat frequency (= flight tone) and wing length is far more contentious however, with conflicting reports on potential correlations (see e.g. [26, 27]). We measured wing lengths for each group and found significant differences between the sexes (Two-way ANOVA; p<0.001; Figure 1c). Further differences were found between *dsxF*^*+/-*^ and *dsxF*^*+/+*^, as well as *dsxF*^*-/-*^ and *dsxF*^*+/+*^, mosquitoes of both sexes (Two-way ANOVA; p<0.001). Individual correlation analyses for each group showed a relationship between wing length and wing beat frequency only for *dsxF*^*+/+*^ females (Supplemental Figure 1). Furthermore, a linear model fit including data from all groups found no significant relationship between wing length and wing beat frequency (see Supplemental Table 1).

### Male, but not female, mosquitoes show positive phonotactic responses to acoustic stimuli mimicking female flight tones

We tested for mosquito responses to auditory stimulation in order to investigate whether mutants showed altered behaviour (Figure 2a). No females from any genotype showed a significantly greater response to an acoustic stimulus (defined as an approach to the sound source) than to silence (ANOVA; p>0.05 for all comparisons; see Figure 2b top). All male groups tested were found to respond more strongly to tones of 400Hz, the stimulus which most closely mimicked wildtype female WBF, than any other stimulus type (ANOVA; p<0.05; Table 2; Figure 2b bottom). However, a few males also responded to the 100 and 700Hz tones. It seems noteworthy that the males’ flight-mediated responses to the playback tones were equally strong in mutants and controls, suggesting that the *dsxF*^*-/-*^ allele does not affect male flight behaviour.

**Table 2:** Quantification of phonotactic responses to acoustic stimulation. (Top) Median values of the number of responders to coarse phonotactic stimulation for dsxF^+/+^, dsxF^+/-^ and dsxF^-/-^ XX and XY mosquitoes, with SEM values provided in brackets. Significant differences found within a genotype between the response to 400Hz and 100/700Hz stimulation are starred (ANOVA; *p < 0.05). (Bottom) Median values of the number of responders to focused phonotactic stimulation for dsxF^+/+^ XY mosquitoes, with SEM values provided in brackets. Significant differences found between the three stimulation frequencies are starred (ANOVA; ***p < 0.001).

| Sample size (coarse) | 3 cages of 25 | 2 cages of 25 | 3 cages of 25 | 3 cages of 25 | 3 cages of 25 | 3 cages of 25 |
| --- | --- | --- | --- | --- | --- | --- |
| Proportion of responders to control | 0.02 (0.02) | 0.1 (0) | 0 (0) | 0 (0) | 0 (0) | 0 (0.01) |
| Proportion of responders to 100 Hz | 0 (0.03) | 0.09 (0.01) | 0 (0.01) | 0.1* (0.03) | 0.15* (0.03) | 0.1* (0.08) |
| Proportion of responders to 400 Hz | 0.05 (0.02) | 0.11 (0.04) | 0 (0) | 0.69 (0.18) | 0.85 (0.07) | 0.55 (0.17) |
| Proportion of responders to 700 Hz | 0.05 (0.03) | 0.09 (0.04) | 0 (0.02) | 0* (0.01) | 0* (0) | 0.05* (0.02) |
| Sample size (focused) | - | - | - | 6 cages of 25 | - | - |
| Proportion of responders to 380 Hz | - | - | - | 0.75 (0.03) | - | - |
| Proportion of responders to 432 Hz | - | - | - | 0.54*** (0.03) | - | - |
| Proportion of responders to 497 Hz | - | - | - | 0.20*** (0.02) | - | - |

We then investigated if the flight tone differences observed between females with different allelic combinations of *dsxF* (^*+/+, +/-, -/-*^) were behaviourally relevant. Specifically, we tested the phonotactic preferences of *dsxF*^*+/+*^ males to pure tones with frequencies equivalent to the median frequencies of females from all three genotypes (^*+/+*^ = 380 ± 30.0 Hz, ^*+/-*^ = 432 ± 28.7 Hz and ^*+/-*^ = 497 ± 22.2 Hz), at the narrow temperature range of 21 - 22 °C. Males were found to respond significantly more to tones similar to ‘wildtype’ *dsxF*^*+/+*^ female flight tones than to tones mimicking either of the female mutants. The ability of flight tones to induce male phonotaxis followed a ‘dose-dependent’ pattern with *dsxF*^*+/+*^ > *dsxF*^*+/-*^ > *dsxF*^*-/-*^ (ANOVA; p<0.001; Table 2; Figure 2c).

## Discussion

Hearing plays a crucial role in mosquito copulation [28]. The phonotactic responses of mosquito males to the flight tones of nearby flying females (or to artificial pure tones mimicking such females), are an important behavioural feature for mosquito reproductive fitness, and reproduction [29]. As such, the ‘acoustic fitness’ of transgenic lines marked for release in the field is a key requirement for the successful spread of deleterious mutations into wildtype populations. Here we show that the transgenic disruption of a female-specific isoform of the sex-determination gene *doublesex* (*dsxF*) changes female flight tones and that mutant flight tones elicit substantially reduced phonotactic responses in control males. The flight tone changes observed were more pronounced in homozygous (*dsxF*^*-/-*^) than in heterozygous condition (*dsxF*^*-/-*^), indicating a *dsxF*^*+*^ dose-dependence of this phenotypic trait, which contrasts with the previously shown haplosufficiencies [10].

Previous recordings of mosquito flight tones have implemented a variety of analytic techniques, but rarely implemented strict environmental controls. This is problematic given the significant variation for reported flight tones at different temperatures, and also the suggested correlations of flight tone and body size [25, 28]. Here we strictly controlled temperature, and also measured wing length as a proxy for body size, to control for this variability. *Anopheles* swarms form predominantly at dusk, when both light and temperature decrease rapidly [30]. It seems possible that during this time female *Anopheles* flight tones decrease rapidly in direct correlation to these temperature decreases; female *Aedes aegypti* WBF fell by around 10Hz per degree over similar temperature changes [25]. Given the sizeable differences we observed in male phonotactic responses to acoustic stimuli less than 50Hz apart, these differences could have a significant effect on male auditory behaviours.

If *dsxF* mutants are to be released in the wild, then only heterozygous males are likely to be released. Updated cage trials with a starting allelic frequency of only 2.5% predicted population collapse within 14 generations [31]. Generation of *dsx* mutants in other mosquito species (such as *Aedes aegypti*) could not only provide a promising control method to combat other mosquito populations, but also provide an ideal tool to investigate the fundamental mechanisms which underly the sizeable sexual dimorphisms in mosquito auditory systems and behaviours. The *dsxF* isoform is reported to be female-specific, it is therefore reassuring that we found no differences in flight tones between male genotypes. All males not only displayed typical phonotactic behaviour but furthermore retained their acoustic preference for the flight tones of control (‘wildtype’) females around 400Hz (at 20-21°C). Most interestingly also, the intersex phenotype of *dsxF*^*-/-*^ females did not include the display of phonotactic behaviour, possibly indicating an independence of male phonotaxis from the *dsx* pathway or leaving a role for the male *doublesex* isoform (*dsxM*).

Lab-based assays in cage conditions can only partially, at best, replicate field conditions. Throughout our phonotaxis experiments, we provided only a single, monofrequent acoustic stimulus at any one time. This is a poor simulation of the auditory landscape of an *An. gambiae* swarm containing many hundreds of males whose flight tones may be constantly modulated [25]. The presence of multiple females within this environment may lead to selection choices for individual males. This may exacerbate the phonotactic preferences we discovered (see Figure 2c), with males possibly disregarding the flight tones of mutant females if simultaneously presented with the sounds of wildtype ones.

Yet the fact that mutant males retain a strong preference for the sounds of wildtype females, bodes well for the effective spread of mutant alleles into resident wildtype populations. It remains to be seen though whether mutant males can successfully join the natural swarms, in which *Anopheles* copulation occurs. Further studies of both mutant and wildtype swarming behaviour are necessary to better understand – and predict – the relevant male-female interactions. This also holds true for a potential female choice element; although we here found no differences between the male genotypes in terms of flight tones, there may be other differences which influence female mate selection.

The argument for utilising transgenic *Anopheles* strains for fighting malaria grows stronger with each new report of insecticide resistance or change in biting behaviour. It is essential however that transgenic lines are tested thoroughly for their suitability. Not only will such experimental testing improve a respective line’s chances of success, but it will also help to create a more detailed profile of the specific requirements for successful release lines (e.g. gene drive carriers). Given the importance of audition for all disease-transmitting mosquito species, acoustic (and auditory) fitness will feature high on that list of requirements. Acoustic courtship in *Anopheles*, finally, is inextricably linked to the mating swarm. Including swarming behaviour in the pre-release testing will thus be crucial. A pipeline of testing focused on mosquito acoustic mating behaviour could significantly help in boosting the efficacy of any release effort. This testing could comprise a sequence of analyses, covering anatomical investigation of the ear, functional tests of hearing, flight tone recordings, and phonotaxis/ mating assays under cage or semi-field conditions. This study utilised only a fraction of these analyses and discovered ecologically relevant differences between mutant and control lines; a comprehensive assessment may provide substantially more evidence which can inform the decision-making process over mutant release strategies and help optimise future disease-control efforts.

## Acknowledgements

The authors would like to thank Carla Siniscalchi (Imperial College London) for providing G3 strain pupae. This work received funding through a pump-priming award from the BBSRC Vector Borne Disease (VBD) Network ANTI-VeC (AV/PP/0028/1, to J.T.A. and M.S.) and a UCL Global Challenges Research Fund (GCRF) small grant (to J.T.A.) and the European Research Council (ERC) under the Horizon 2020 research and innovation programme (Grant agreement No 648709, to J.T.A.).

## Author contributions

M.P.S., M.G., A.C. and J.T.A. contributed to the conception and design of the research. M.P.S., M.G., K.K. and J.B. performed experiments. M.P.S., M.G. and J.T.A. analysed the data. M.P.S., M.G. and J.T.A. wrote the manuscript. J.T.A supervised the study. All authors read and approved of the final manuscript.

## Ethics approval and consent to participate

Not applicable.

## Consent for publication

Not applicable.

### Competing interests

The authors declare no competing interests.

## Data availability

All data analysed in this paper are available from the authors, as well as more comprehensive details on experimental or analytical methodologies.

## Figure Legends

**Supplementary Figure 1:**
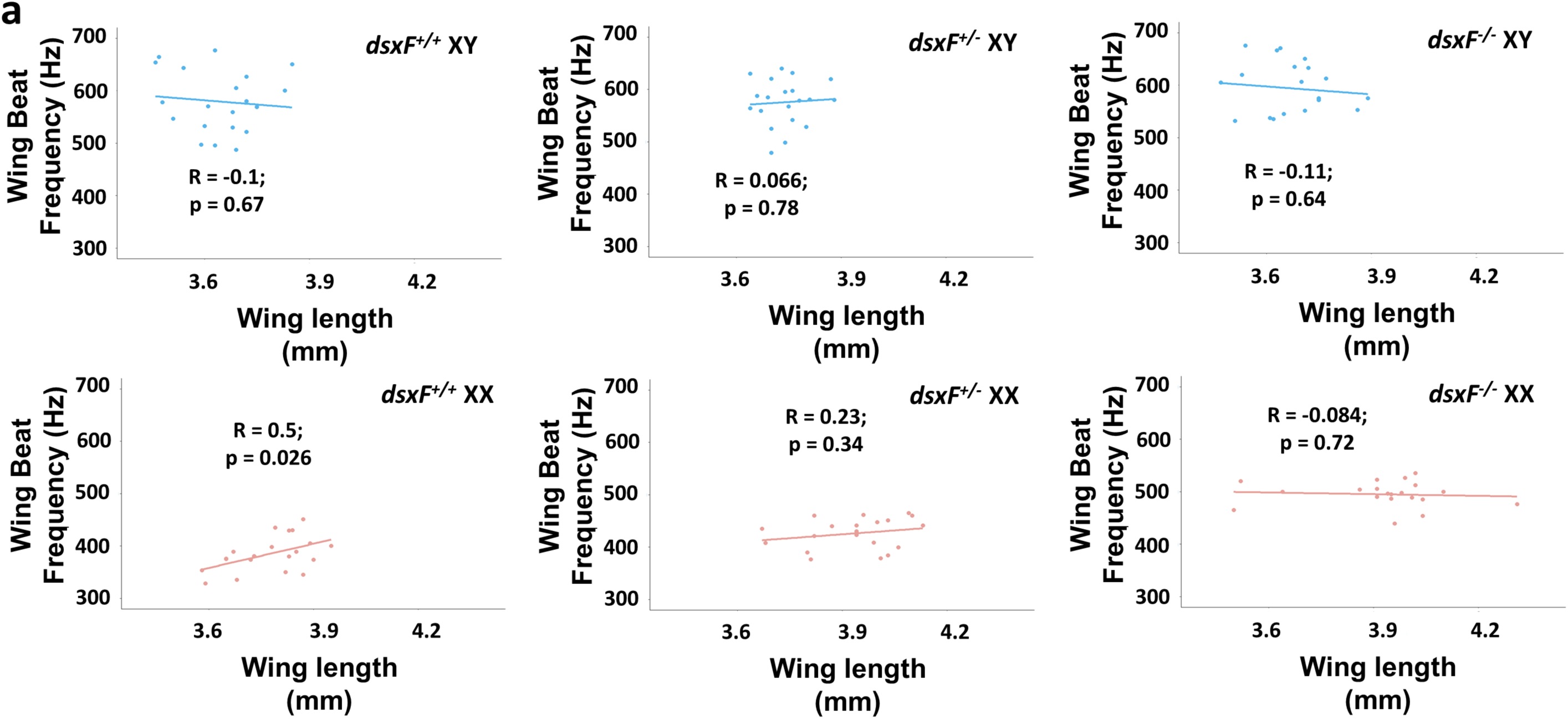
Correlations between wing length and wing beat frequency. a) Correlations between wing length (mm) and wing beat frequency (Hz) for all groups tested. Sample sizes are the same as for wing beat frequency calculations.

**Supplementary Figure 2:**
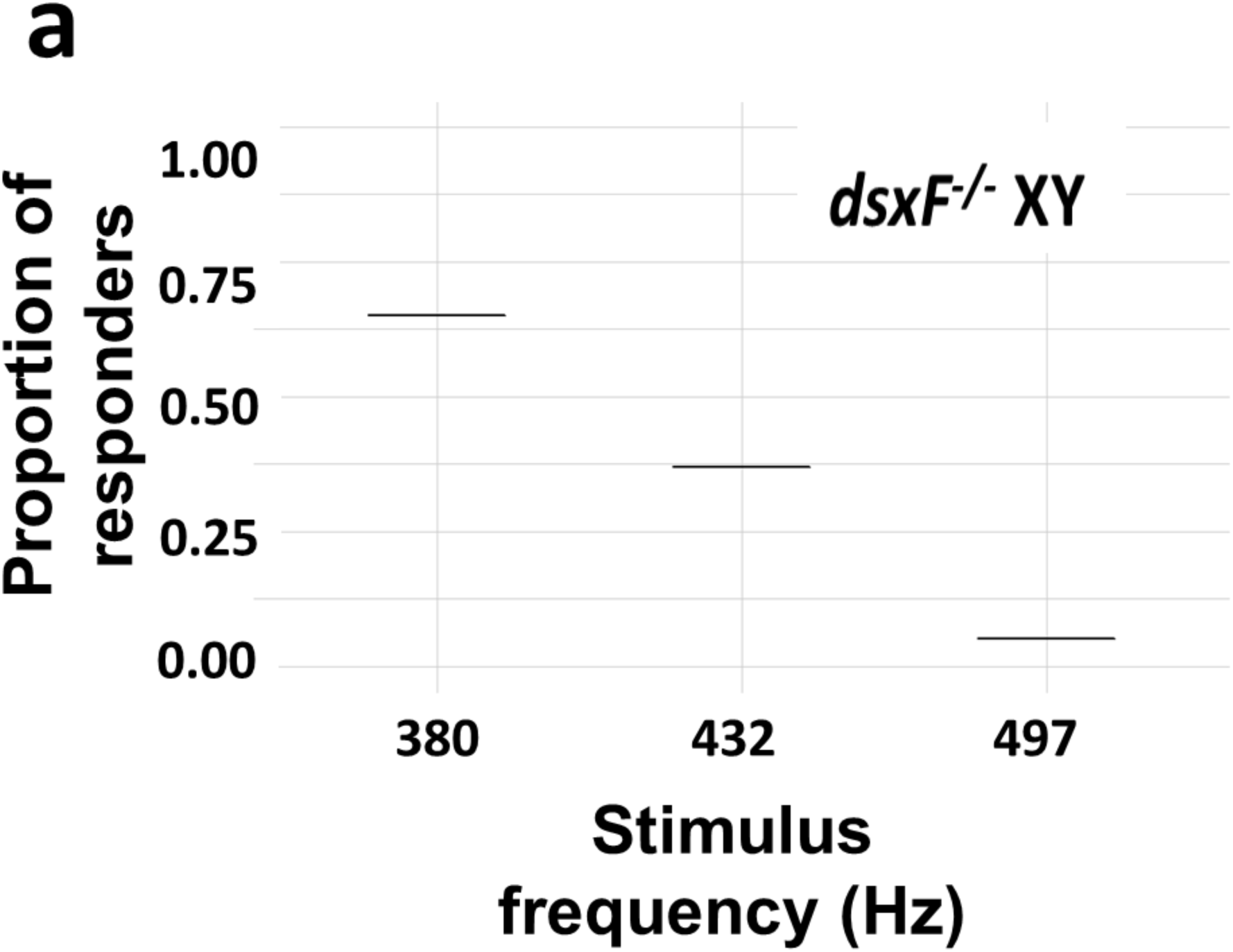
Phonotactic response of *dsxF*^*-/-*^ males to phonotactic stimulation. a) Adjusted proportion of control mosquitoes responding to each stimulus type (380Hz, 432Hz and 497Hz respectively) for *dsxF*^*-/-*^ XY mosquitoes. Centre line, median; error bars represent ± SEM.

## Supplemental Information

### Linear model fitting

In order to investigate the potential relationship between wing beat frequency (=flight tones) and wing length (as well as other potential variables) in greater detail, we used the R package ‘lme4’ to fit the following equation:

Wing beat frequency ∼ Sex*Genotype + Wing length

We found that sex, genotype, and sex:genotype were all highly significant factors in determining wing beat frequency. However, wing length was not found to significantly affect wing beat frequency.

---

**Supplemental table 1.**
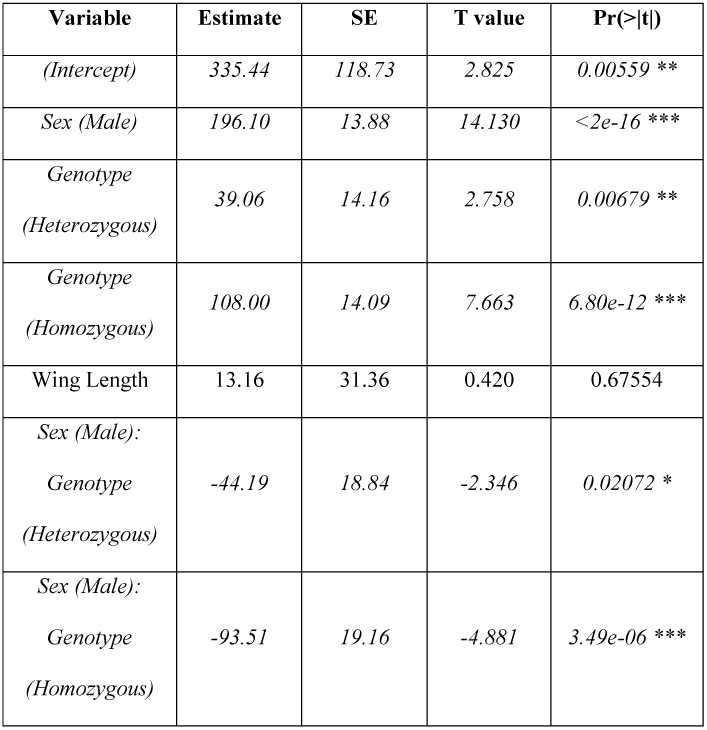
Outputs of linear model relating wing beat frequency to sex, genotype and wing length. Significant values are italicised.

## Notes

### Competing Interest Statement

The authors have declared no competing interest.

## References

1. James SL, Abate D, Abate KH, Abay SM, Abbafati C, Abbasi N, et al. Global, regional, and national incidence, prevalence, and years lived with disability for 354 diseases and injuries for 195 countries and territories, 1990-2017: a systematic analysis for the Global Burden of Disease Study 2017. Lancet. 2018;392 10159:1789–858.

2. O’Meara WP, Mangeni JN, Steketee R, Greenwood B. Changes in the burden of malaria in subSaharan Africa. Lancet Infect Dis. 2010;10 8:545–55.

3. WHO: Malaria Fact Sheet (https://www.who.int/en/news-room/fact-sheets/detail/malaria).https://www.who.int/en/news-room/fact-sheets/detail/malaria (2019).

4. Ranson H, N’Guessan R, Lines J, Moiroux N, Nkuni Z, Corbel V. Pyrethroid resistance in African anopheline mosquitoes: what are the implications for malaria control? Trends Parasitol. 2011;27 2:91–8.

5. Lwetoijera DW, Harris C, Kiware SS, Dongus S, Devine GJ, McCall PJ, et al. Increasing role of Anopheles funestus and Anopheles arabiensis in malaria transmission in the Kilombero Valley, Tanzania. Malaria J. 2014;13.

6. Sougoufara S, Ottih EC, Tripet F. The need for new vector control approaches targeting outdoor biting anopheline malaria vector communities. Parasite Vector. 2020;13 1:295.

7. Hemingway J, Shretta R, Wells TNC, Bell D, Djimde AA, Achee N, et al. Tools and Strategies for Malaria Control and Elimination: What Do We Need to Achieve a Grand Convergence in Malaria? Plos Biol. 2016;14 3.

8. Deredec A, Godfray HCJ, Burt A. Requirements for effective malaria control with homing endonuclease genes. Proceedings of the National Academy of Sciences of the United States of America. 2011;108 43:E874–E80.

9. Flores HA, O’Neill SL. Controlling vector-borne diseases by releasing modified mosquitoes. Nat Rev Microbiol. 2018;16 8:508–18.

10. Kyrou K, Hammond AM, Galizi R, Kranjc N, Burt A, Beaghton AK, et al. A CRISPR-Cas9 gene drive targeting doublesex causes complete population suppression in caged Anopheles gambiae mosquitoes. Nature Biotechnology. 2018;36 11:1062-+.

11. Champer J, Buchman A, Akbari OS. Cheating evolution: engineering gene drives to manipulate the fate of wild populations. Nat Rev Genet. 2016;17 3:146–59.

12. Brossard D, Belluck P, Gould F, Wirz CD. Promises and perils of gene drives: Navigating the communication of complex, post-normal science. Proceedings of the National Academy of Sciences of the United States of America. 2019;116 16:7692–7.

13. Aldersley A, Pongsiri A, Bunmee K, Kijchalao U, Chittham W, Fansiri T, et al. Too “sexy” for the field? Paired measures of laboratory and semi-field performance highlight variability in the apparent mating fitness of Aedes aegypti transgenic strains. Parasite Vector. 2019;12.

14. Leftwich PT, Bolton M, Chapman T. Evolutionary biology and genetic techniques for insect control. Evol Appl. 2016;9 1:212–30.

15. Soma DD, Maiga H, Mamai W, Bimbile-Somda NS, Venter N, Ali AB, et al. Does mosquito mass-rearing produce an inferior mosquito? Malaria J. 2017;16.

16. Pennetier C, Warren B, Dabire KR, Russell IJ, Gibson G. “Singing on the Wing” as a Mechanism for Species Recognition in the Malarial Mosquito Anopheles gambiae. Curr Biol. 2010;20 2:131–6.

17. Clements A. THE BIOLOGY OF MOSQUITOES VOLUME 3 TRANSMISSION OF VIRUSES AND INTERACTIONS WITH BACTERIA Preface. Biology of Mosquitoes, Vol 3: Transmission of Viruses and Interactions with Bacteria. 2012:Viii-+.

18. Aldersley A, Cator LJ. Female resistance and harmonic convergence influence male mating success in Aedes aegypti. Sci Rep-Uk. 2019;9.

19. Manoukis NC, Diabate A, Abdoulaye A, Diallo M, Dao A, Yaro AS, et al. Structure and Dynamics of Male Swarms of Anopheles gambiae. J Med Entomol. 2009;46 2:227–35.

20. Belton P. Attraction of Male Mosquitos to Sound. J Am Mosquito Contr. 1994;10 2:297–301.

21. Albert Joerg T, Kozlov Andrei S. Comparative Aspects of Hearing in Vertebrates and Insects with Antennal Ears. Curr Biol. 2016;26 20:R1050–R61.

22. Andres M, Seifert M, Spalthoff C, Warren B, Weiss L, Giraldo D, et al. Auditory Efferent System Modulates Mosquito Hearing. Curr Biol. 2016;26 15:2028–36.

23. Su MP, Andrés M, Boyd-Gibbins N, Somers J, Albert JT. Sex and species specific hearing mechanisms in mosquito flagellar ears. Nature Communications. 2018;9 1:3911.

24. Fraenkel G. Untersuchungen über die Koordination von Reflexen und automatisch-nervösen Rhythmen bei Insekten. Zeitschrift für vergleichende Physiologie. 1932;16 2:371–93.

25. Villarreal SM, Winokur O, Harrington L. The Impact of Temperature and Body Size on Fundamental Flight Tone Variation in the Mosquito Vector Aedes aegypti (Diptera: Culicidae): Implications for Acoustic Lures. J Med Entomol. 2017;54 5:1116–21.

26. Cator LJ, Ng’Habi KR, Hoy RR, Harrington LC. Sizing up a mate: variation in production and response to acoustic signals in Anopheles gambiae. Behavioral Ecology. 2010;21 5:1033–9.

27. Pantoja-Sanchez H, Gomez S, Velez V, Avila FW, Alfonso-Parra C. Precopulatory acoustic interactions of the New World malaria vector Anopheles albimanus (Diptera: Culicidae). Parasite Vector. 2019;12.

28. Andrés M, Su MP, Albert J, Cator LJ. Buzzkill: targeting the mosquito auditory system. Current Opinion in Insect Science. 2020;40:11–7.

29. Stone CM, Tuten HC, Dobson SL. Determinants of Male Aedes aegypti and Aedes polynesiensis (Diptera: Culicidae) Response to Sound: Efficacy and Considerations for Use of Sound Traps in the Field. J Med Entomol. 2013;50 4:723–30.

30. Sawadogo SP, Costantini C, Pennetier C, Diabaté A, Gibson G, Dabiré RK. Differences in timing of mating swarms in sympatric populations of Anopheles coluzzii and Anopheles gambiae s.s. (formerly An. gambiae M and S molecular forms) in Burkina Faso, West Africa. Parasite Vector. 2013;6 1:275.

31. Simoni A, Hammond AM, Beaghton AK, Galizi R, Taxiarchi C, Kyrou K, et al. A male-biased sex-distorter gene drive for the human malaria vector Anopheles gambiae. Nature Biotechnology. 2020.

